# Integrated characterization of SARS-CoV-2 genome, microbiome, antibiotic resistance and host response from single throat swabs

**DOI:** 10.1101/2020.10.15.340794

**Authors:** Bo Lu, Yi Yan, Liting Dong, Lingling Han, Yawei Liu, Junping Yu, Jianjun Chen, Danyang Yi, Meiling Zhang, Chao Wang, Runkun Wang, Dengpeng Wang, Hongping Wei, Di Liu, Chengqi Yi

## Abstract

The ongoing coronavirus disease 2019 (COVID-19) pandemic, caused by severe acute respiratory syndrome coronavirus 2 (SARS-CoV-2) infection, poses a severe threat to humanity. Rapid and comprehensive analysis of both pathogen and host sequencing data is critical to track infection and inform therapies. In this study, we performed unbiased metatranscriptomic analysis of clinical samples from COVID-19 patients using a newly-developed RNA-seq library construction method (TRACE-seq), which utilizes tagmentation activity of Tn5 on RNA/DNA hybrids. This approach avoids the laborious and time-consuming steps in traditional RNA-seq procedure, and hence is fast, sensitive and convenient. We demonstrated that TRACE-seq allowed integrated characterization of full genome information of SARS-CoV-2, putative pathogens causing coinfection, antibiotic resistance and host response from single throat swabs. We believe that the integrated information will deepen our understanding of pathogenesis and improve diagnostic accuracy for infectious diseases.

## Introduction

Longstanding, emerging, and re-emerging infectious diseases continuously threaten human health across centuries(1). Precise and rapid identification of pathogens from clinical samples is important for both guiding infection treatment strategies and monitoring novel infectious disease outbreaks, e.g. the outbreak of SARS-CoV-2, in the community. While most nucleic acid amplification-based and pathogen specific antibody detection-based molecular techniques only detect a limited number of pathogens and need their prior knowledge, metagenomic or meta-transcriptomic approaches allow for comprehensive and unbiased identification and characterization of microbiome directly from clinical specimens(2).

Compared to meta-genomic sequencing, meta-transcriptomic sequencing has several distinct advantages: it permits detection of RNA viruses that would not be interpreted in metagenomic data, reveals transcriptionally active organism(s) which are more etiologically important, and indicates host immune response which is essential to distinguish true pathogens from colonizers(3-5). However, the laborious and time-consuming steps in traditional RNA-seq experiments hinder the development of meta-transcriptomic based clinical diagnostics for rapid pathogen identification.

Very recently, we and others have independently developed a rapid and cost-effective RNA-seq method, based on Tn5 tagmentation activity towards RNA/DNA hybrids(6, 7). Our method, termed “TRACE-seq”, enables rapid one-tube library construction for RNA-seq experiments and shows excellent performance in compsrison to traditional RNA-seq methods. We thus envisioned that this convenient and sensitive method could be applied to clinical specimens for unbiased meta-transcriptomic analysis. In this study, we modified the TRACE-seq procedure, shorten the total time and optimized analytical pipeline to meet the needs for clinical meta-transcriptomic diagnosis and analysis. We then applyed TRACE-seq to meta-transcriptomic sequencing of single throat swabs specimens from COVID-19 patients and healthy individuals. We found library construction of specimens could be accomplished in ∼2h with high quality. Analysis of TRACE-seq meta-transcriptomic data of 13 SARS-CoV-2 positive samples and 2 negative samples demonstrated the success of this method to sensitively detect SARS-CoV-2 with high coverage even for samples with relatively high Ct values, or to assemble unknown microbe genome *de novo* (using SARS-CoV-2 as an example). Moreover, TRACE-seq sensitively detected the microbiome and simultaneously allowed for interrogating antibiotic resistance and host responses. Taken together, TRACE-seq enables unbiased pathogen detection and could have broad applications in meta-transcriptomic study and clinical diagnosis.

## Results

### TRACE-seq enables metatranscriptomic analysis

To perform metatranscriptomic analysis on clinical samples, such as throat swabs in this study, we made several modifications to TRACE-seq. First, to achieve unbiased sequencing of microbiome, we used both random hexamer and oligo d(T)_23_VN primers for reverse transcription, using approximately 1/10 total RNA extracted from a single throat swab as input. Secondly, we reduced the total time of library construction to around 2 hours (Figure 1a), which enables TRACE-seq to be more compatible for clinical use, especially when substantial numbers of specimens require investigation. Third, we developed a tailored analytical pipeline of TRACE-seq to simultaneously identify known and unknown pathogens and at the meanwhile to characterize host transcriptional response in a single metatranscriptomic profiling reaction (Figure 1b). This new pipeline allowed us to obtain rich information from the metatranscriptomic data generated by the modified TRACE-seq.

**Figure 1.**
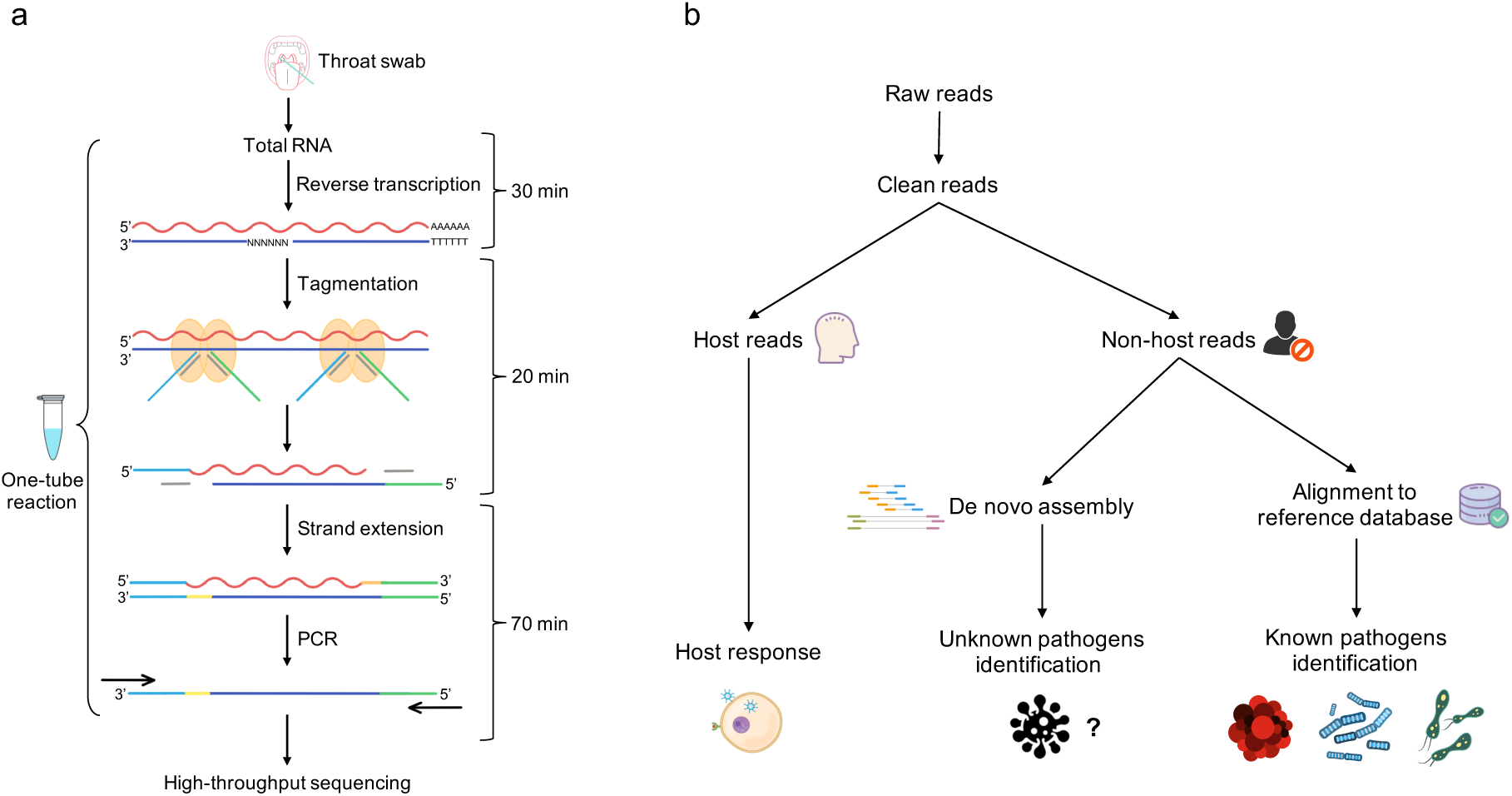
Workflow of TRACE-seq enabled metatranscriptomic sequencing for clinical diagnosis. **a**. A wet lab protocol of TRACE-seq starting with total RNA extracted from throat swabs of COVID-19 patients. **b**. A dry lab pipeline including known and unknown pathogens identification and host response characterization.

### Sensitive detection of SARS-CoV-2 genome

Since the samples were positive or suspected positive throat swabs from COVID-19 patients, we asked whether the untargeted meta-transcriptomic sequencing could yield a full genome sequence of SARS-CoV-2 virus. After removing low quality reads and human reads, the remaining reads were mapped to the SARS-CoV-2 reference genome Wuhan-Hu-1 (accession number: NC_045512). Sequencing covered the reference genome from 171 bp to 29,903 bp (0.57%-100%), with an average sequencing depth from 2.56× to 44,737× (Supplementary table 1). Four samples (B101, A193, B13, C1) with low Ct values showed poor coverage and sequencing depth; the isolated RNA samples from the throat swabs were repeatedly frozen and thawed, probably compromising the integrity of RNA and hence were excluded. The other nine samples were used for subsequent correlation analysis. Among the remaining nine samples, the proportion of obtained reads of SARS-CoV-2, the coverage to the reference genome, the average sequencing depth and the median sequencing depth all showed a negative correlation with the Ct value of the samples (Spearman test, p < 0.05) (Figure 2a). In addition, the whole genome sequence could be acquired from mapping-based approach when the Ct value is as high as 32 (n=4, 44.44% of samples), with the average sequencing depth of 131×. Even in samples with Ct values up to 35 (n=7, 77.78% of samples), more than 94% of genome can be covered by TRACE-seq (Figure 2b, Supplementary table 1, Supplementary figure 1).

**Figure 2.**
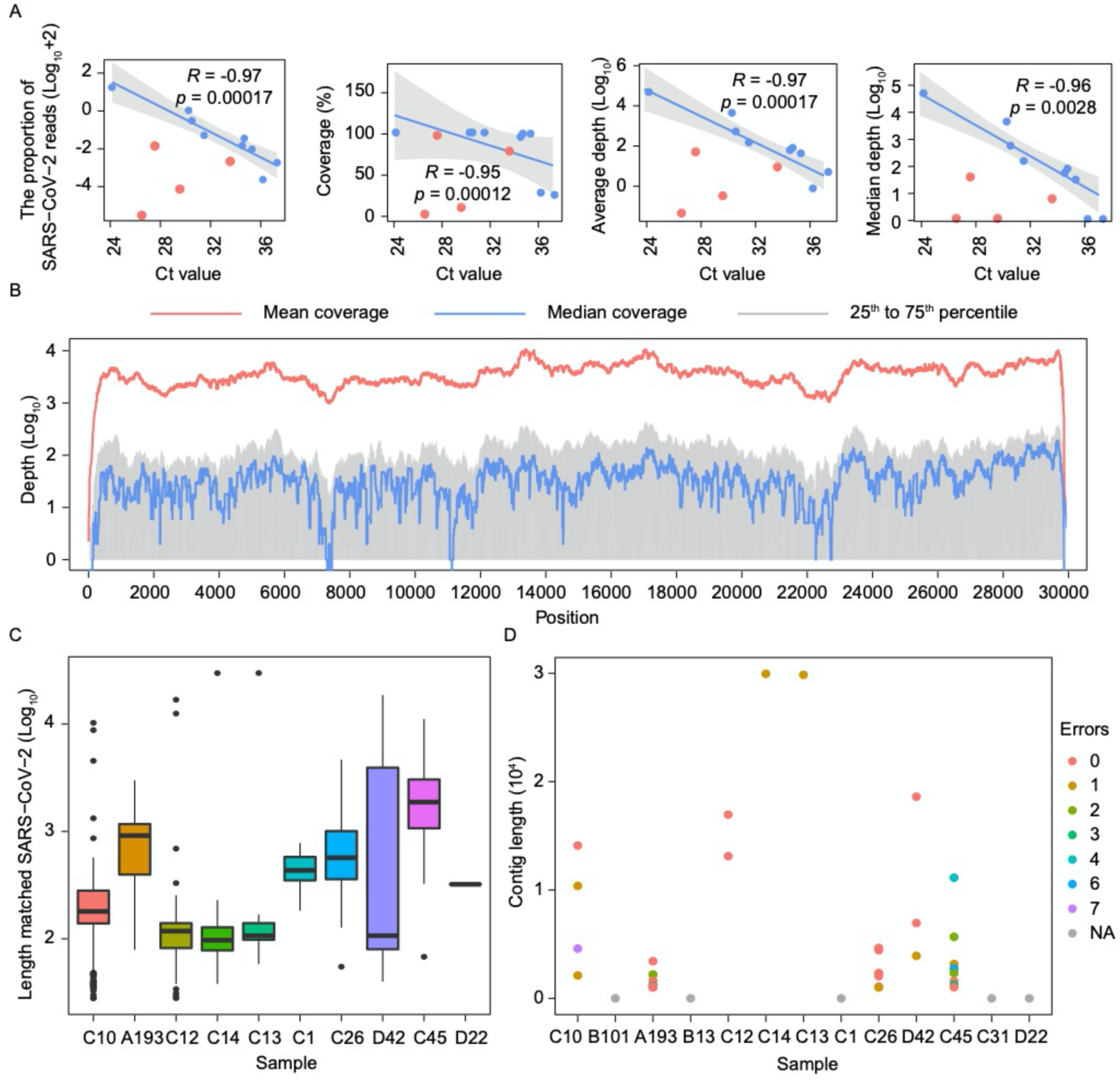
Genome coverage of SARS-CoV-2. **a**. Correlation between SARS-CoV-2 sequencing reads and Ct value in 13 positive samples. From the left to the right: the correlation of the ratio of SARS-CoV-2 reads, the coverage of SARS-CoV-2 genome, the average sequencing depth, the median sequencing depth and the Ct value of each sample are shown in order. The red dots represent samples with abnormal sequencing results, and linear regression indicates the relationship between the sequencing data and the Ct value of samples with normal sequencing results (blue dots). **b**. Genome coverage of sequenced samples across the SARS-CoV-2 genome. The x axis represents the virus genome position, y axis represents the log10 depth of each site. Lines in red represent the mean sequencing depth, lines in blue represent the median sequencing depth, and areas in grey represent 25^th^ to 75^th^ percentile of sequencing depth. **c**. Length distribution of contigs matched SARS-CoV-2. The x axis represents each sample, and the y axis represents log_10_ lengths of contigs matched SARS-CoV-2. **d**. De novo assembly results of SARS-CoV-2. The graph shows contigs only when the length of matched to the SARS-COV-2 genome over 1,000 bp. The y axis represents length of contigs of each sample (the x axis). Different colors represent the number of error bases (shown in legends) in each contig relative to previously known genome sequences.

### Reconstruction of full-length genome of SARS-CoV-2

Of the 452,865 contigs (average 34,836, from 18,160 to 53,415) assembled *de-novo* from non-human reads, 3,500 contigs (average 269, from 0 to 2,976) were determined to be SARS-CoV-2 genome fragments. There were no SARS-CoV-2 contigs in sample C31, B101 and B13. Most of contigs (n=3,461, 98.89%) were less than 1,000 bp (Figure 2c). To determine the accuracy of this method in acquisition of pathogen genome, all SARS-CoV-2 contigs were searched against genomes of each sample. In contigs with matched length over 1,000 bp, most contigs (22/39, 56.41%) were completely consistent with their corresponding genome (Figure 2d), while the other contigs had error bases from 1 to 7. In samples C14 and C13 with excellent coverage and depth, almost full-length genome (29,793 bp and 29,825 bp) were obtained just from *de-novo* assembly. Thus, TRACE-seq could enable the *de-novo* assembly of the complete genome of unknown pathogens and be readily utilized to identify emerging pathogens in patients with unknown etiology of infection and efficiently complement routine diagnostics.

### Unbiased identification of putative pathogens in addition to SARS-CoV-2

It is widely reported that coinfection (multi-species infection) contributes to enhanced morbidity and mortality, especially in elderly and immunosuppressed influenza patients(8, 9). Thus, we were curious to see if our metatranscriptomic sequencing approach could capture other pathogens in addition to SARS-CoV-2. Indeed, alignment of TRACE-seq data to microbe reference databases identified many bacteria, fungi and viruses in both patient and healthy samples (Figure 3a). To assess whether COVID-19 patients and healthy individuals have different microbe community in their throat, principal coordinates analysis (PCoA) was conducted using relative abundance of the microbiome. We observed that COVID-19 patients harbored a throat microbiome that is quite different from healthy individuals (Figure 3b). In addition, sample C31 differed significantly from other SARS-CoV-2 positive samples. Further investigation of the relative abundance of probable respiratory pathogens revealed that patient C31 contained the most abundant *Klebsiella pneumoniae* and Human gammaherpesvirus 4, compared to other samples (Figure 3c), which might be a cause of the separation between sample C31 and the rest of the SARS-CoV-2 positive samples.

**Figure 3.**
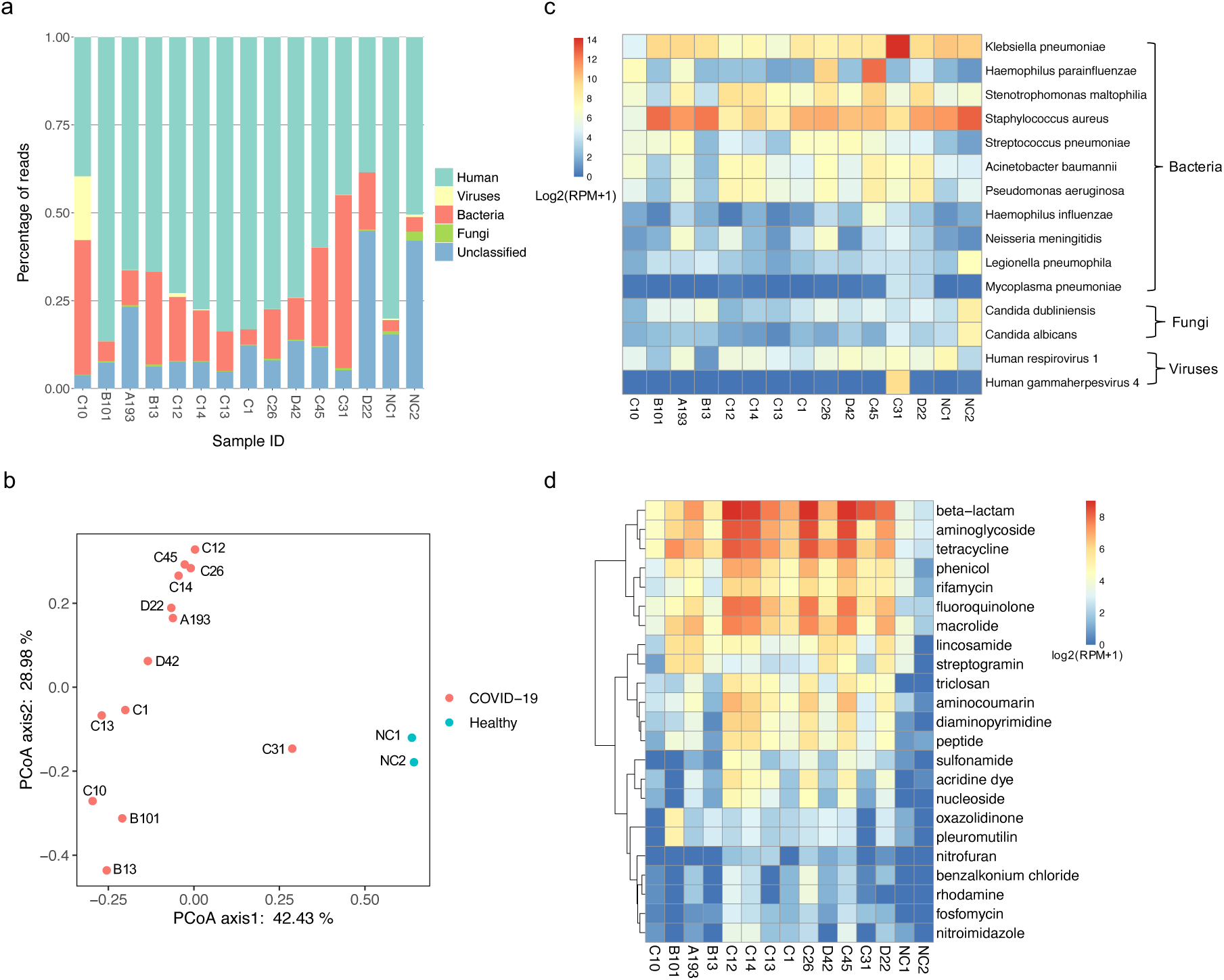
Microbiome profiles in COVID-19 patients and healthy individuals. **a**. Histogram showing percentage of reads mapping to human, viruses, bacteria and fungi for the individual samples. **b**. PCoA of microbiome using relative abundance at the genus level. **c**. Heatmap showing relative abundance of potential respiratory pathogens identified in SARS-CoV-2 positive and negative samples. RPM: reads per million non-host reads. **d**. Heatmap displaying relative abundance of antibiotic resistance genes in SARS-CoV-2 positive and negative samples.

Among the probable respiratory pathogens listed in Figure 3c, *Stenotrophomonas maltophilia,Haemophilus parainfluenzae, Staphylococcus aureus, Streptococcus pneumoniae, Haemophilus influenzae* and *Acinetobacter baumannii* are common commensal organism of the normal oropharynx, however, they can also become opportunistic pathogens and cause infectious disease, such as endocarditis, bacteremia and pneumonia(10-13). *Klebsiella pneumoniae, Stenotrophomonas maltophilia, Pseudomonas aeruginosa, Neisseria meningitidis* and *Legionella pneumophila* cause disease infrequently in normal hosts but can be a major cause of infection in patients with underlying or immunocompromising conditions(14-18). *Mycoplasma pneumoniae* is a type of “atypical” bacteria that commonly causes mild infections of the respiratory system(19). As for identified fungi, *Candida dubliniensis* and *Candida albicans* are both opportunistic yeast and can be detected in the gastrointestinal tract in healthy adults; they were also known to cause respiratory diseases (20-22). Human respirovirus 1, also known as Human parainfluenza virus 1, is the most common cause of croup and also associated with pneumonia. Human gammaherpesvirus 4 is one of the most common viruses in human; it is best known as the cause of infectious mononucleosis (23, 24), and is also constantly detected in lungs of patients with idiopathic pulmonary fibrosis (25). In our results, a relatively high abundance of *Haemophilus parainfluenzae, Streptococcus pneumoniae, Acinetobacter baumannii, Pseudomonas aeruginosa* and *Neisseria meningitidis* were identified in several SARS-CoV-2 positive samples compared with negative samples, which indicated potential coinfection. Nevertheless, these data by itself could not prove that COVID-19 patients were coinfected by these identified microorganism; these data have to be carefully interpreted in the clinical context.

### Profiles of antibiotic resistance genes

Antimicrobial resistance has become a global issue. Pathogens with antibiotic resistance are increasing and many pathogens are becoming multidrug-resistant (26, 27). To characterize antibiotic resistance gene expression profiles, we aligned metatranscriptomic reads against the Comprehensive Antibiotic Resistance Database (CARD) (28). On average, around 84 antibiotic resistance genes were identified in SARS-CoV-2 positive samples, while only around 23 genes were identified in negative samples. According to the CARD, the identified antibiotic resistance genes confer resistance to 23 classes of antibiotics. Almost all resistance gene classes were more abundant in COVID-19 patients compared to healthy individuals. Genes conferring resistance to beta-lactam, aminoglycoside, tetracycline, phenicol, rifamycin, fluoroquinolone and macrolide were the most abundant (Figure 3d). Overall, the distinct microbiome, emergence of potential coinfection, and the elevated abundance of antibiotic resistance genes provide new data for establishing clinical therapeutic scheme during the treatment for COVID-19 patients.

### Characterization of host response to SARS-CoV-2

Distinguishing infection from colonization remains challenging. Because host transcriptional profiling has emerged as a promising diagnostic tool for infectious diseases (29, 30), we next tested whether the host response to SARS-CoV-2 could be simultaneously characterized by TRACE-seq mediated metatranscriptomic analysis from throat swabs. As shown in Figure 3a, a substantial percentage of the reads are derived from human, and an average of 14,766 human genes with FPKM > 1 were detected per sample (Figure 4a, Figure S2a and b). Based on the gene expression profiles, the relationships between samples were inspected using a multidimensional scaling (MDS) plot (Figure 4b). As expected, SARS-CoV-2 positive samples were clearly separated from negative samples. In addition, sample C31 differed significantly in host gene expression from other SARS-CoV-2 positive samples, which might be caused by the relatively high abundance of Klebsiella pneumoniae and Human gammaherpesvirus 4 identified in sample C31. To characterize the common host response to SARS-CoV-2, we excluded sample C31 when performing differential gene expression analysis between SARS-CoV-2 positive and negative samples. We identified 153 differentially expressed genes, 149 of which were up-regulated (Figure 4c, Figure S2c). Gene Ontology enrichment analysis identified the top up-regulated biological processes to be immune response, defense response, viral process and response to cytokine (Figure 4d). Further investigation revealed that a subset of up-regulated genes involve in IL1B-associated inflammatory response (IL1B, IL8, IL36A, CXCR2, FOS, ANXA1, CASP4, KRT16, S100A8, S100A9). Moreover, another subset of up-regulated genes (ISG15, EGR1, IFI27, IFIT2, IFIT3, IFITM1, IFITM2, IFITM3, HLA-B, HLA-C) were enriched in type I interferon signaling pathway (Figure 4e). These results were highly consistent with previously reported host response to SARS-CoV-2 (31-33). Overall, metatranscriptomic data via TRACE-seq of throat swab samples demonstrates reliable performance in characterization of host transcriptional response to the infection of SARS-CoV-2.

**Figure 4.**
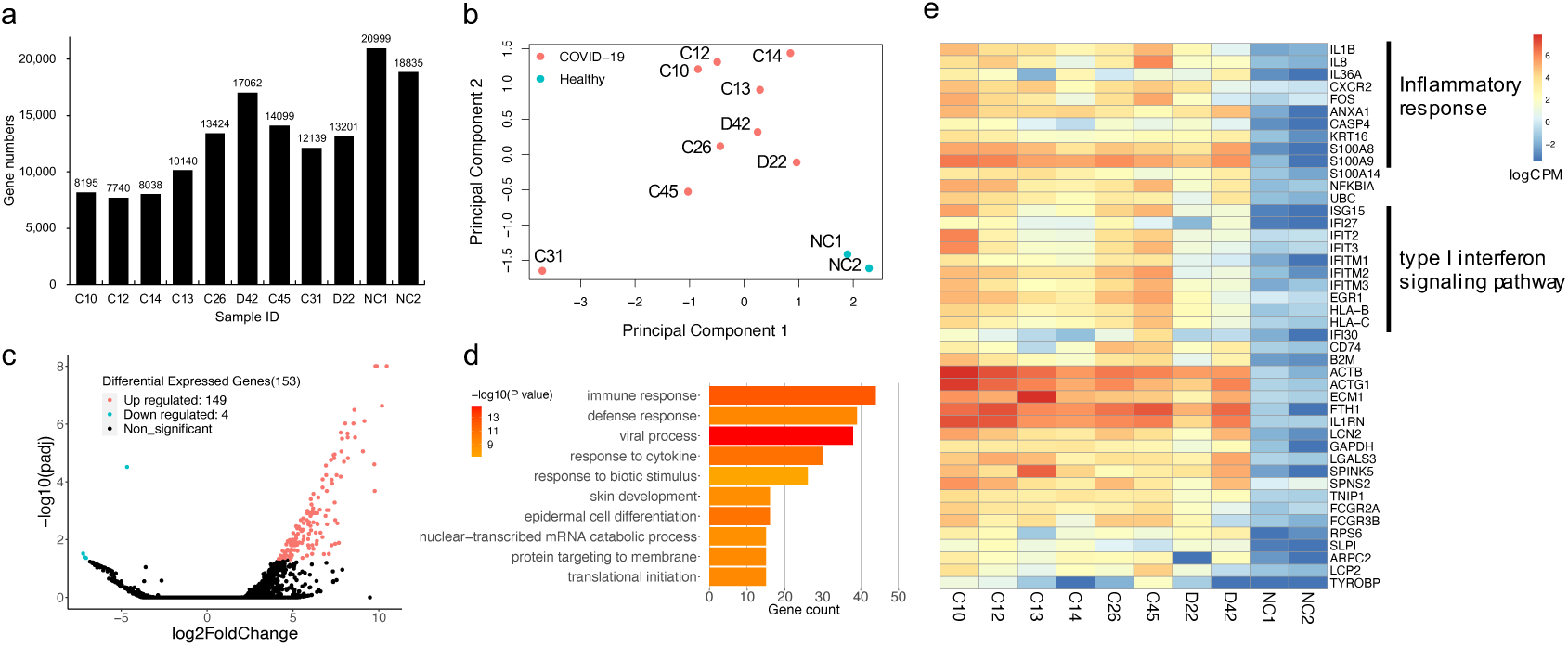
Profiling of host transcriptional response. **a**. Bar plot showing gene numbers detected in each sample. **b**. MDS plot showing variation among samples based on host transcriptional profiles. **c**. Volcano plot showing differentially expressed genes between SARS-CoV-2 positive and negative samples. Significantly up- and down-regulated genes (padj < 0.05, |log2FoldChage| > 1) are highlighted in red and blue, respectively. **d**. Bar plot of the most enriched Gene Ontology terms. **e**. Heatmap presenting the differentially expressed immune response related genes between SARS-CoV-2 positive and negative samples.

## Discussion

Although next generation sequencing holds a great potential to directly detect known and unknown pathogens including viruses, bacteria, fungi and parasites in a single application, the laborious and time-consuming steps in traditional RNA library construction procedure hinders its clinical application. As a rapid and convenient one-tube RNA-seq library construction method, TRACE-seq significantly lower the barrier for extensive application of unbiased RNA-seq in clinical diagnosis. In addition, multiplexing libraries by utilizing Tn5 transposase containing barcoded adaptors could enable sample investigation in a high-throughput manner, particularly when comprehensive surveillance for emerging pathogens is needed during a sudden disease outbreak.

It is very challenging to discriminate pathogens from background commensal microbiota, since substantive bacteria or fungi can colonize multiple body sites of healthy individuals. The microbe present at a relatively higher abundance in patients compared to healthy individuals are often considered as a pathogen, yet the abundance thresholds indicating infection is difficult to define based solely on microbiome information. On the other hand, host transcriptional profiling has been reported to distinguish infectious and noninfectious diseases (30) and to further discriminate between virial and bacterial infections (29). A previous study integrates host response and unbiased microbe detection for lower respiratory tract infection diagnosis in critically ill adults, using both RNA-seq and DNA-seq but yet lacking antibiotic resistance analysis (3). Another study characterized microbial gene expression profiles (including antibiotic resistance genes) using nasal and throat swab samples, and host response using blood samples during influenza infection (34). To our knowledge, this is the first study integrating unbiased pathogens detection, antibiotic resistance and host response in a single approach with throat swabs from COVID-19 patients. In our results, SARS-CoV-2 positive and negative samples differed significantly in both microbiome composition and host response. Among SARS-CoV-2 positive samples, sample C31 harbored a throat microbiome and host response notably distinct from others, indicating sufficiently different pathogens present in patient C31 compared to other samples. Moreover, TRACE-seq hold the potential to construct a network of microbiome composition, antibiotic resistance and host response for characterizing the complex host-microbiome interactions. Ideally, TRACE-seq data can be utilized to develop a model combining pathogens metric, antibiotic resistance and host transcriptional classifier for infectious diseases diagnosis. We believe that the integrated information acquired from a TRACE-seq library will deepen our understanding of pathogenesis, improve diagnostic accuracy and more precisely inform optimal antimicrobial treatment for infectious diseases caused by not only SARS-CoV-2 but also other pathogens and eventually facilitate the utility of metatranscriptomic profiling as a routine diagnostic method.

## Materials and methods

### Ethics statement

The study and use of all samples were approved by the Ethics Committee of Wuhan Institue of Virology (No. WIVH17202001).

### Sample collection and nucleotide extraction

Respiratory specimens (swabs) collected from patients admitted to various Wuhan health care facilities were immediately placed into sterile tubes containing 3 ml of viral transport media (VTM). The swabs were deactivated by heating at 56°C for 30 minutes in a biosafety level 2 (BSL 2) laboratory at the Wuhan Institute of Virology in Zhengdian Park with personal protection equipment for biosafety level 3 (BSL 3) laboratory. Total nucleic acids were extracted using QIAamp 96 virus Qiacube HT kit on QIAxtractor Automated extraction (Qiagen, US) following the manufacturer’s instructions.

### TRACE-seq library preparation and sequencing

TRACE-seq libraries were constructed using TruePrep® RNA Library Prep Kit for Illumina (Vazyme, TR502-01) according to the manufacturer’s instructions with several modifications. 1/10 volume of total nucleic acids extracted from each swab was used for each library. After 18 PCR cycles, the library was purified using 0.8X Agencourt AMPure XP beads (Beckman Coulter) and eluted in 20 μl nuclease-free water. The concentration of resulting libraries was determined by Qubit 3.0 fluorometer with the Qubit dsDNA HS Assay kit (Invitrogen) and the size distribution of libraries was assessed by Agilent 2100 Bioanalyzer. Finally, libraries were sequenced on the Illumina Hiseq X10 platform which generated 2 x 150 bp of paired-end raw reads.

### Data preprocessing

Raw reads from sequencing were firstly subjected to Trim Galore (v0.6.4_dev) (http://www.bioinformatics.babraham.ac.uk/projects/trim_galore/) for quality control and adaptor trimming. The minimal threshold of quality was 20, and the minimal length of reads to remain was set as 20 nt.

### Host transcriptional profiling analysis

Filtered reads were mapped to human genome (hg19) and transcriptome using STAR (v2.7.1a) (35). The FPKM value for annotated genes was calculated by cuffnorm (v2.2.1) (36), and genes with FPKM > 1 were considered to be expressed. Multidimensional scaling and differential gene expression analysis were conducted using EdgeR (v3.28.1) (37) with gene count data generated by HTSeq (v0.11.2) (38). Gene Ontology Enrichment Analysis for biological processes was performed by DAVID (v6.8) (39) with all significantly up-regulated genes as input. Due to the redundancy of enriched GO terms, GO terms and their p values were further summarized using REViGO (40). The top 10 enriched representative GO terms were plotted.

### Discrimination and de-novo assembly of SARS-CoV-2

After removal of human reads, the remaining data were aligned to the reference genome of Wuhan-Hu-1 (GenBank accession number: NC_045512) using Bowtie2 (v2.2.9) (41) for SARS-CoV-2 identification. The coverage and sequencing depth of SARS-CoV-2 genome were calculated by Samtools (v1.9) (42). On the other hand, to verify the method could screen for aetiologic agents and obtain pathogen genome, all non-human reads were processed for de-novo assembly using MEGAHIT (v1.2.9) with default parameters (43), and then all contigs were searched against NCBI nt database using blastn for classification(44). As for accuracy of assembly sequences, contigs determined to come from SARS-CoV-2 were performed blastn (with the parameter “-outfmt 3”) to display the differences with corresponding genome.

### Microbiome analysis

After removing human reads, the remaining reads were subjected to microbial taxonomic classification using Kraken2 (v2.0.8-beta) (45) with a custom database. To build the custom database, standard RefSeq complete bacterial genomes were downloaded through “kraken2-build --download-library bacteria” and complete genomes of human viruses and genome assemblies of fungi were downloaded from NCBI’s RefSeq and added to the custom database’s genomic library using the “--add-to-library” switch. Principal coordinate analysis (PCoA) of relative abundances of microbial taxa at the genus level was done using cmdscale command in R. Distances between samples were calculated using Morisita-horn dissimilarity index by vegdist command from vegan package version 2.5-6 (https://CRAN.R-project.org/package=vegan). The antibiotic resistance genes were annotated by aligning the filtered metatranscriptomic reads to the Comprehensive Antibiotic Resistance Database (CARD). Antibiotic resistance genes with more than 10 completely matching reads were considered. The relative expression of antibiotic resistance genes were determined as RPM (reads per million non-host reads). All corresponding graphs were plotted using R scripts by RStudio (v1.2.5033) (https://rstudio.com/).

## Acknowledgments

The authors would like to thank Vazyme Biotech in Nanjing, China, for assisting in library procedure optimization and providing library preparation kits. In addition, the authors would like to thank National Center for Protein Sciences at Peking University in Beijing, China, for assistance with experiments. Part of the analysis was performed on the High Performance Computing Platform of the Center for Life Science (Peking University). This work was supported by International Innovation Resource Cooperation Project, Beijing Municipal Science and Technology Commission (No. to C.Y.), National Natural Science Foundation of China (nos. 31861143026, 91740112 and 21825701 to C.Y.) and Epidemic Prevention and Control Special Project, Peking University.

## Contributions

C.Q.Y. and D.L. conceived the project; C.Q.Y., D.L., H.P.W., and D.P.W. supervised the project; B.L., Y.Y., and L.T.D. designed the experiments together and wrote the manuscript; L.L.H. performed experiments with the help of C.W. and R.W.; B.L. and Y.Y. performed the bioinformatics analysis; J.P.Y. collected the clinical samples; J.J.C., D.Y.Y., M.L.Z., and Y.W.L. participated in discussion.

## References

1. Lozano R, Naghavi M, Foreman K, Lim S, Shibuya K, Aboyans V, et al. Global and regional mortality from 235 causes of death for 20 age groups in 1990 and 2010: a systematic analysis for the Global Burden of Disease Study 2010. Lancet. 2012;380(9859):2095–128.

2. Chiu CY, Miller SA. Clinical metagenomics. Nature reviews Genetics. 2019;20(6):341–55.

3. Langelier C, Kalantar KL, Moazed F, Wilson MR, Crawford ED, Deiss T, et al. Integrating host response and unbiased microbe detection for lower respiratory tract infection diagnosis in critically ill adults. Proceedings of the National Academy of Sciences of the United States of America. 2018;115(52):E12353–E62.

4. Langelier C, Zinter MS, Kalantar K, Yanik GA, Christenson S, O’Donovan B, et al. Metagenomic Sequencing Detects Respiratory Pathogens in Hematopoietic Cellular Transplant Patients. Am J Respir Crit Care Med. 2018;197(4):524–8.

5. Gliddon HD, Herberg JA, Levin M, Kaforou M. Genome-wide host RNA signatures of infectious diseases: discovery and clinical translation. Immunology. 2018;153(2):171–8.

6. Lu B, Dong L, Yi D, Zhang M, Zhu C, Li X, et al. Transposase-assisted tagmentation of RNA/DNA hybrid duplexes. Elife. 2020;9.

7. Di L, Fu Y, Sun Y, Li J, Liu L, Yao J, et al. RNA sequencing by direct tagmentation of RNA/DNA hybrids. Proceedings of the National Academy of Sciences of the United States of America. 2020;117(6):2886–93.

8. Fischer N, Indenbirken D, Meyer T, Lutgehetmann M, Lellek H, Spohn M, et al. Evaluation of Unbiased Next-Generation Sequencing of RNA (RNA-seq) as a Diagnostic Method in Influenza Virus-Positive Respiratory Samples. J Clin Microbiol. 2015;53(7):2238–50.

9. Chertow DS, Memoli MJ. Bacterial coinfection in influenza: a grand rounds review. JAMA. 2013;309(3):275–82.

10. Man WH, de Steenhuijsen Piters WA, Bogaert D. The microbiota of the respiratory tract: gatekeeper to respiratory health. Nat Rev Microbiol. 2017;15(5):259–70.

11. Mitchell JL, Hill SL. Immune response to Haemophilus parainfluenzae in patients with chronic obstructive lung disease. Clin Diagn Lab Immunol. 2000;7(1):25–30.

12. Smeltzer MS. Staphylococcus aureus Pathogenesis: The Importance of Reduced Cytotoxicity. Trends Microbiol. 2016;24(9):681–2.

13. Howard A, O’Donoghue M, Feeney A, Sleator RD. Acinetobacter baumannii: an emerging opportunistic pathogen. Virulence. 2012;3(3):243–50.

14. Paczosa MK, Mecsas J. Klebsiella pneumoniae: Going on the Offense with a Strong Defense. Microbiol Mol Biol Rev. 2016;80(3):629–61.

15. Brooke JS. Stenotrophomonas maltophilia: an emerging global opportunistic pathogen. Clin Microbiol Rev. 2012;25(1):2–41.

16. Sadikot RT, Blackwell TS, Christman JW, Prince AS. Pathogen-host interactions in Pseudomonas aeruginosa pneumonia. Am J Respir Crit Care Med. 2005;171(11):1209–23.

17. Overturf GD. Indications for the immunological evaluation of patients with meningitis. Clin Infect Dis. 2003;36(2):189–94.

18. Kumpers P, Tiede A, Kirschner P, Girke J, Ganser A, Peest D. Legionnaires’ disease in immunocompromised patients: a case report of Legionella longbeachae pneumonia and review of the literature. J Med Microbiol. 2008;57(Pt 3):384–7.

19. Kashyap S, Sarkar M. Mycoplasma pneumonia: Clinical features and management. Lung India. 2010;27(2):75–85.

20. AbdulWahab A, Salah H, Chandra P, Taj-Aldeen SJ. Persistence of Candida dubliniensis and lung function in patients with cystic fibrosis. BMC Res Notes. 2017;10(1):326.

21. Shweihat Y, Perry J, 3rd, Shah D. Isolated Candida infection of the lung. Respir Med Case Rep. 2015;16:18–9.

22. Wahab AA, Taj-Aldeen SJ, Kolecka A, ElGindi M, Finkel JS, Boekhout T. High prevalence of Candida dubliniensis in lower respiratory tract secretions from cystic fibrosis patients may be related to increased adherence properties. Int J Infect Dis. 2014;24:14–9.

23. Stanfield BA, Luftig MA. Recent advances in understanding Epstein-Barr virus. F1000Res. 2017;6:386.

24. Dunmire SK, Hogquist KA, Balfour HH. Infectious Mononucleosis. Curr Top Microbiol Immunol. 2015;390(Pt 1):211–40.

25. Tang YW, Johnson JE, Browning PJ, Cruz-Gervis RA, Davis A, Graham BS, et al. Herpesvirus DNA is consistently detected in lungs of patients with idiopathic pulmonary fibrosis. J Clin Microbiol. 2003;41(6):2633–40.

26. Boolchandani M, D’Souza AW, Dantas G. Sequencing-based methods and resources to study antimicrobial resistance. Nature reviews Genetics. 2019;20(6):356–70.

27. Tillotson GS, Zinner SH. Burden of antimicrobial resistance in an era of decreasing susceptibility. Expert Rev Anti Infect Ther. 2017;15(7):663–76.

28. McArthur AG, Waglechner N, Nizam F, Yan A, Azad MA, Baylay AJ, et al. The comprehensive antibiotic resistance database. Antimicrob Agents Chemother. 2013;57(7):3348–57.

29. Suarez NM, Bunsow E, Falsey AR, Walsh EE, Mejias A, Ramilo O. Superiority of transcriptional profiling over procalcitonin for distinguishing bacterial from viral lower respiratory tract infections in hospitalized adults. J Infect Dis. 2015;212(2):213–22.

30. Tsalik EL, Henao R, Nichols M, Burke T, Ko ER, McClain MT, et al. Host gene expression classifiers diagnose acute respiratory illness etiology. Sci Transl Med. 2016;8(322):322ra11.

31. Lee JS, Park S, Jeong HW, Ahn JY, Choi SJ, Lee H, et al. Immunophenotyping of COVID-19 and influenza highlights the role of type I interferons in development of severe COVID-19. Sci Immunol. 2020;5(49).

32. Zhou Z, Ren L, Zhang L, Zhong J, Xiao Y, Jia Z, et al. Heightened Innate Immune Responses in the Respiratory Tract of COVID-19 Patients. Cell Host Microbe. 2020;27(6):883–90 e2.

33. Ong EZ, Chan YFZ, Leong WY, Lee NMY, Kalimuddin S, Haja Mohideen SM, et al. A Dynamic Immune Response Shapes COVID-19 Progression. Cell Host Microbe. 2020;27(6):879–82 e2.

34. Zhang L, Forst CV, Gordon A, Gussin G, Geber AB, Fernandez PJ, et al. Characterization of antibiotic resistance and host-microbiome interactions in the human upper respiratory tract during influenza infection. Microbiome. 2020;8(1):39.

35. Dobin A, Davis CA, Schlesinger F, Drenkow J, Zaleski C, Jha S, et al. STAR: ultrafast universal RNA-seq aligner. Bioinformatics. 2013;29(1):15–21.

36. Trapnell C, Williams BA, Pertea G, Mortazavi A, Kwan G, van Baren MJ, et al. Transcript assembly and quantification by RNA-Seq reveals unannotated transcripts and isoform switching during cell differentiation. Nat Biotechnol. 2010;28(5):511–5.

37. Robinson MD, McCarthy DJ, Smyth GK. edgeR: a Bioconductor package for differential expression analysis of digital gene expression data. Bioinformatics. 2010;26(1):139–40.

38. Anders S, Pyl PT, Huber W. HTSeq--a Python framework to work with high-throughput sequencing data. Bioinformatics. 2015;31(2):166–9.

39. Huang da W, Sherman BT, Lempicki RA. Systematic and integrative analysis of large gene lists using DAVID bioinformatics resources. Nat Protoc. 2009;4(1):44–57.

40. Supek F, Bosnjak M, Skunca N, Smuc T. REVIGO summarizes and visualizes long lists of gene ontology terms. PLoS One. 2011;6(7):e21800.

41. Langmead B, Salzberg SL. Fast gapped-read alignment with Bowtie 2. Nature methods. 2012;9(4):357–9.

42. Li H, Handsaker B, Wysoker A, Fennell T, Ruan J, Homer N, et al. The Sequence Alignment/Map format and SAMtools. Bioinformatics. 2009;25(16):2078–9.

43. Li D, Liu CM, Luo R, Sadakane K, Lam TW. MEGAHIT: an ultra-fast single-node solution for large and complex metagenomics assembly via succinct de Bruijn graph. Bioinformatics. 2015;31(10):1674–6.

44. Camacho C, Coulouris G, Avagyan V, Ma N, Papadopoulos J, Bealer K, et al. BLAST+: architecture and applications. BMC Bioinformatics. 2009;10:421.

45. Wood DE, Lu J, Langmead B. Improved metagenomic analysis with Kraken 2. Genome Biol. 2019;20(1):257.

